# Daytime Napping and Associative-Memory Retrieval Speed: An Exploratory Behavioral and EEG Study

**DOI:** 10.64898/2026.10.19.760160

**Authors:** Minqi Yang, Ruohan Wang, Yawen Guan, Huimin Zhao, Weicong Wang, Liangdong Ji, Yiling Liu, Linxi Yan, Chunyu Qu

## Abstract

**Purpose:** To test whether a daytime nap was associated with improved associative-recognition performance relative to monitored wakefulness and to determine where any response-time (RT) difference occurred in the RT distribution.

**Methods:** Fifteen healthy students completed nap and wake sessions in a within-participant design. Each session included word-pair learning, immediate recognition, a 60-min interval, and delayed recognition without additional study. Trial-level log RT was the focal outcome; accuracy and electroencephalography (EEG) were complementary outcomes.

**Results:** The RT Condition × Time interaction was significant (*b* = −0.063, 95% *CI* [−0.117, −0.009], *p* = .027). Responses were 6.7% faster after the nap (*p* = .010) but did not change after wakefulness (*p* = .704), with the difference concentrated among slower responses. Accuracy showed no condition-specific change (*p* = .401). Nap EEG showed aperiodic spectral slowing and stronger periodic sigma activity, but no EEG measure explained the RT effect.

**Conclusions:** Daytime napping was associated with faster delayed associative recognition, primarily through attenuation of the slowest responses. The exploratory findings warrant larger confirmatory studies.

## Introduction

Paired-associate tasks assess the retention of relations between studied elements and are widely used to examine declarative memory across a post-learning interval. Improvement at a later test can reflect several processes, including offline stabilization, retrieval practice, changes in alertness, and resistance to interference; by itself, it does not establish a specific consolidation mechanism [1, 2]. Daytime naps nevertheless provide a practical model for testing whether a period containing sleep is associated with different retention from an equivalent period of wakefulness. Previous studies have reported nap-related benefits for paired-associate retention and resistance to interference [3–7], but other brief-nap studies have found no advantage [8]. This variability is plausible because nap duration and sleep architecture differ markedly between individuals, while initial acquisition, task difficulty, delay length, and the wake comparison activity can all moderate the observable outcome [3, 4, 9, 10].

Most work in this area has emphasized accuracy or retention scores. These measures are central to memory performance, but they can be insensitive when recognition is already high and the remaining errors provide little room for condition-specific improvement. Response time (RT) on correct trials offers complementary information about how efficiently an association is retrieved. Importantly, an average RT difference can arise in at least two ways: the entire distribution may shift by a similar amount, or the conditions may differ mainly in a subset of unusually slow responses. The second pattern is theoretically informative because prolonged responses may reflect difficult retrieval, weak evidence, response uncertainty, or transient attentional lapses. Quantile analysis cannot identify which process generated a slow response, but it can establish whether a condition effect is broadly distributed or localized to the slow tail. Examining the RT distribution therefore provides a more precise behavioral description than relying on mean RT alone and is especially useful when accuracy is near ceiling.

EEG can characterize how the nap and wake intervals differed physiologically and can test whether those differences covary with subsequent behavior. Neurophysiological accounts of sleep-dependent declarative memory emphasize coordinated non-rapid eye movement activity, including slow oscillations, spindles, and hippocampal ripples [11–14]. Yet short naps are heterogeneous, and the mere presence of a sleep feature does not show that it produced a behavioral difference. Moreover, conventional band power combines periodic oscillations with the aperiodic, 1/f-like background. Separating these components can distinguish a genuine narrowband change from a broadband change in spectral slope or offset and can consequently refine interpretations of delta, alpha, sigma, and beta activity [15]. EEG recorded during recognition offers a second level of evidence because retrieval has been associated with frontal and posterior old/new effects and with changes in theta, alpha, and beta power [16–18]. A condition-specific change in retrieval EEG would indicate that the post-interval tests engaged measurably different retrieval-stage activity, whereas its absence would argue against a readily detectable retrieval correlate in this sample.

The present exploratory, within-participant study examined associative recognition before and after a 60-min nap opportunity and a matched period of monitored wakefulness. The focal question was whether RT changed differently between conditions; accuracy provided a complementary test of recognition performance. We then used RT quantiles to determine whether any difference reflected general speeding or attenuation of the slowest responses. Finally, sleep architecture, conventional and parameterized spectra, sleep events, and retrieval EEG were examined as secondary physiological outcomes and potential correlates of behavioral change. Because the study and analyses were not preregistered and the sample was small, these analyses were intended to define a focused behavioral and physiological pattern for later confirmation rather than to establish a specific sleep-dependent mechanism.

## Materials and Methods

### Participants and Design

Eighteen participants were recruited; 15 had usable archived data (mean age 21.53 ± 1.60 years, range 18–25; 7 men, 8 women). Eligibility was assessed by interview, medical history, sleep questionnaires [19, 20], chronotype, body-mass index, and mood and anxiety screening [21, 22]. Exclusion criteria included sleep, neurological, psychiatric, or cognitive disorders and prior participation in a similar experiment. Participants avoided alcohol, caffeine, tobacco, and psychoactive substances before testing. The study was approved by the Ethics Committee of the School of Education, Zhengzhou University (Approval No. JYXY20250928), and all participants provided written informed consent. No a priori power analysis was documented. Each participant completed one nap session and one monitored-wake session on separate days; each interval lasted 60 min. Sessions occurred between 12:30 and 15:00. After EEG setup, participants studied 65 semantic word pairs and completed immediate recognition (Fig. 1). Participants then completed the nap or monitored-wake interval. Delayed recognition began 15 min later without additional learning or restudy. Brief prompts or maintenance tasks were administered every 15–20 min during wakefulness; their timing was not preserved in the archived EEG.

**Fig. 1.**
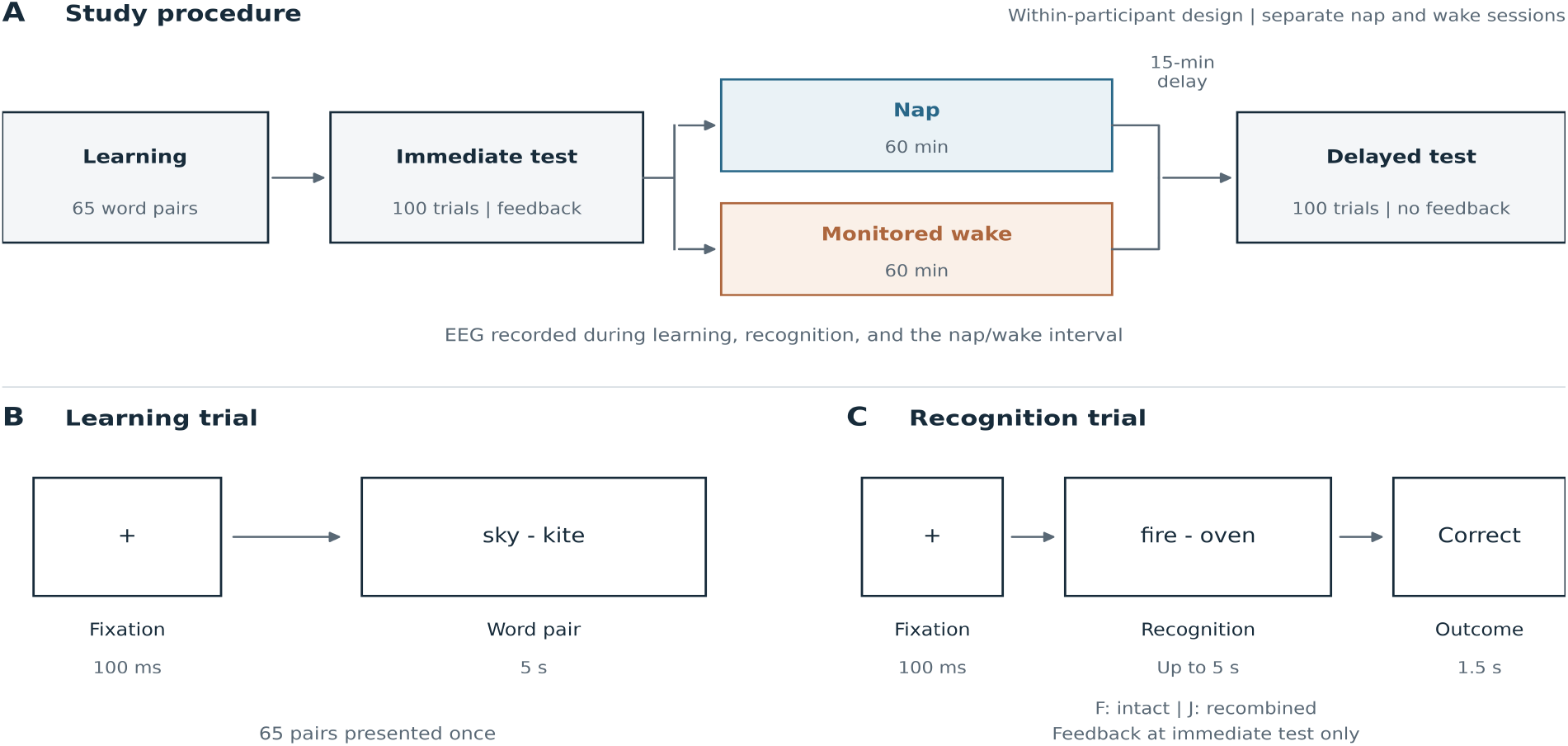
Experimental Design and Task Sequence. (a) Within-participant crossover procedure. (b) Learning trial. (c) Recognition trial

### Associative-Memory Task

Nap and wake sessions used different condition-specific lists; the archived files did not show that lists were exchanged across participants. Each list contained 65 semantically associated study pairs presented for 5 s with a 100-ms fixation interval (Fig. 1). Each recognition test contained 65 intact and 35 recombined pairs. Participants pressed F for intact and J for recombined pairs. Immediate and delayed tests used the same items in randomized orders; the immediate test provided feedback, whereas the delayed test did not.

#### EEG Acquisition and Sleep Scoring

EEG was recorded in separate learning, test, and interval segments using a Neuroscan 64-channel system with EOG and chin EMG. Signals were sampled at 500 Hz and resampled to 200 Hz for EEG analyses; impedances were ≤5 kΩ for EEG/EOG and ≤10 kΩ for EMG. Nap recordings were visually scored in consecutive 30-s epochs according to AASM criteria [23] as wake, N1, N2, N3, or REM, using F3, F4, C3, C4, O1, and O2 referenced to the contralateral mastoid together with EOG and chin EMG, band-pass filtered at 0.3–35 Hz. Epochs obscured by artefact were marked during scoring.

### Behavioral Analysis

Accuracy was the proportion of correct responses. Participant-level mean RT used correct trials after excluding values beyond ±3 SD within each test. Inverse efficiency was mean RT/accuracy, and relative RT improvement was (immediate RT − delayed RT)/immediate RT. Trial-level models were fitted in R with lme4 and lmerTest [24, 25]. ID07’s immediate wake test was missing; the accuracy model included 5,900 trials from 59 tests, and the log-RT model included 5,529 correct trials with RT ≥200 ms. Effect-coded Condition, Time, and their interaction were fixed effects. Final random effects were (1 + Condition × Time | Participant) + (1 | Item) for RT and (1 + Time | Participant) + (1 | Item) for accuracy. RT degrees of freedom used the Satterthwaite method; simple effects used emmeans [26].

Correct-trial RTs were summarized at the 10th, 30th, 50th, 70th, and 90th percentiles within participant, condition, and test. An exact sign-flip test of the linear interaction trend assessed whether the contrast increased toward the slow tail; individual quantiles were exploratory. Sensitivity analyses varied RT processing and examined missing-data and participant influence (Online Resource 1, Tables S1–S3).

### EEG Analysis

EEG was processed in MNE-Python [27]. Data were notch-filtered at 50 Hz, resampled to 200 Hz, cleaned by automated bad-channel detection and interpolation, and decomposed with extended-Infomax ICA; high-probability eye-blink and heartbeat components were removed with ICLabel [28]. Data were band-pass filtered at 0.3–35 Hz and re-referenced to linked mastoids. Recordings were divided into 30-s epochs. Epochs were rejected if scored as artefact, if frontal or central peak-to-peak amplitude exceeded 500 µV, or if 20–35-Hz log power exceeded the recording median by >4 robust SD. Retained data averaged 53.5 ± 6.9 min for nap and 55.3 ± 7.7 min for wake. Sleep measures included time in bed, total sleep time (TST; N1 + N2 + N3 + REM), sleep efficiency, sleep-onset latency, wake after sleep onset, and stage durations.

Power spectral density was estimated for clean epochs with Welch’s method (4-s Hamming windows, 50% overlap, 0.25-Hz resolution) and averaged over frontal (F3, Fz, F4) and central (C3, Cz, C4) regions. Absolute power was integrated over delta (0.5–4 Hz), theta (4–8 Hz), alpha (8–12 Hz), sigma (12–15 Hz), and beta (15–30 Hz) and log10-transformed. Spectra were parameterized with FOOOF over 1–30 Hz (fixed aperiodic mode) [15]. Aperiodic offset and exponent were extracted; periodic power was the residual after removing the aperiodic fit. Mean fit R^2^ was .993 for nap and .971 for wake. Conditions were compared with paired t tests on log10 band power. Holm correction was applied across five bands within each region [29], separately for aperiodic parameters and periodic bands. Processing robustness is reported in Online Resource 1 (Tables S4–S5). Stage-specific spectra, whole-scalp tests, sleep spindles and slow waves (detected with YASA [30]), retrieval EEG, and sleep–behavior associations were secondary analyses (Online Resource 1).

### Data Availability

De-identified derived tables and analysis code are available from the corresponding author on reasonable request. Raw EEG and trial-level files are restricted by the original consent and ethics requirements.

## Results

### Sleep and Behavioral Results

Fifteen participants were included; demographic and sleep characteristics are shown in Table 1.

**Table 1.** Demographic characteristics and sleep habits of the participants (N = 15)

| Variable | *M* | *SD* |
| --- | --- | --- |
| Age (years) | 21.53 | 1.60 |
| Sex, *n* (%) | Men: 7 (46.7%);<br>women: 8 (53.3%) | — |
| Average nap duration per week (min) | 47.50 | 17.80 |
| Number of naps per week | 4.53 | 1.69 |
| PSQI | 2.93 | 1.34 |
| ISI | 4.33 | 1.95 |
| MEQ-5 | 13.00 | 1.36 |

*Note.* Sex is reported as *n* (%); all other variables are reported as *M* and *SD*. PSQI = Pittsburgh Sleep Quality Index; ISI = Insomnia Severity Index; MEQ-5 = reduced Morningness–Eveningness Questionnaire.

Mean TST was 41.1 ± 15.0 min and sleep efficiency was 66.0 ± 23.5% (Table 2). N2 predominated (28.0 ± 16.7 min); four participants showed N3, and no REM sleep was scored.

**Table 2.**
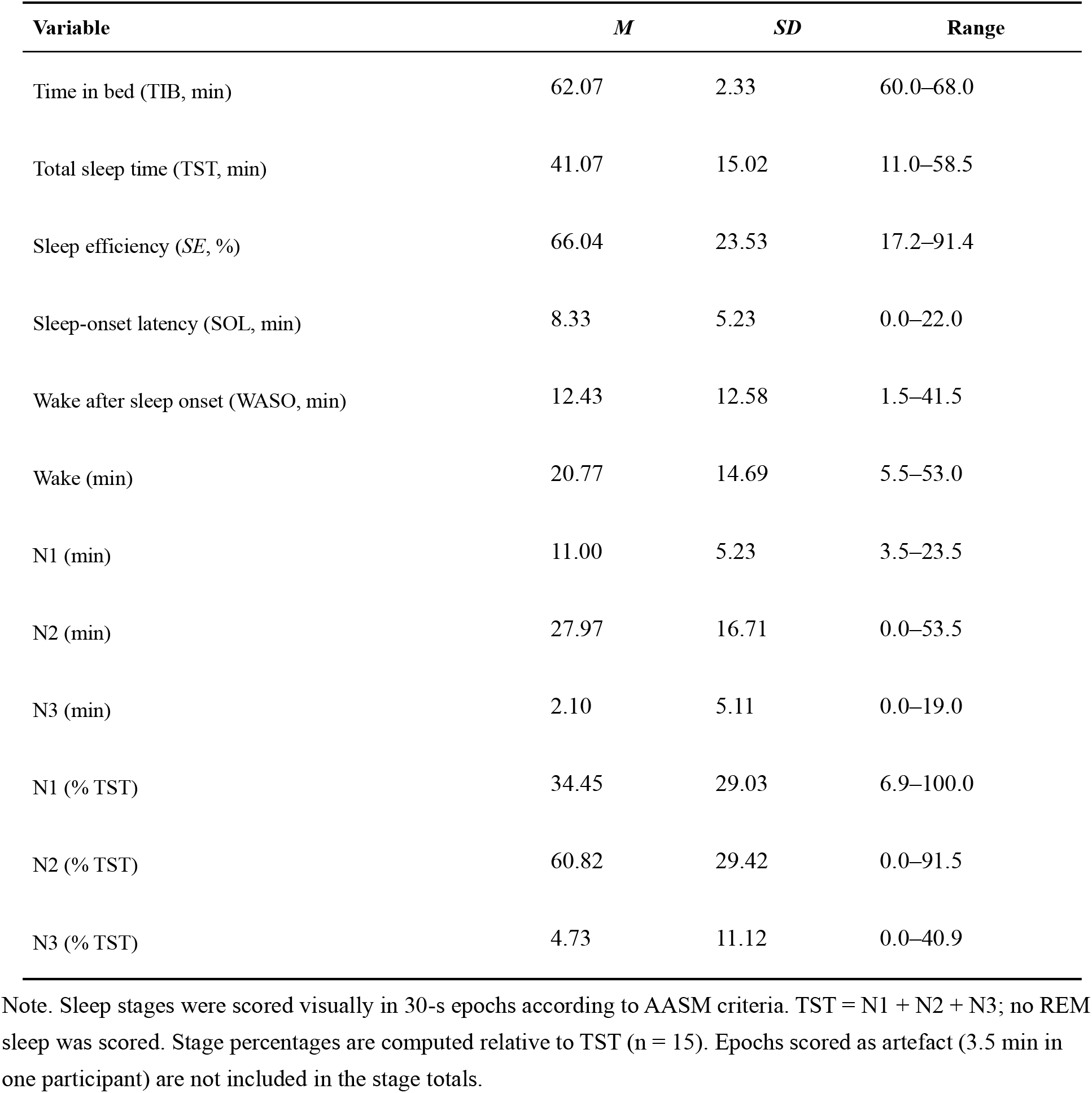
Sleep architecture of the nap (N = 15)

For log RT, the Condition × Time interaction was significant (*b* = −0.063, *SE* = 0.025, 95% *CI* [−0.117, −0.009], *t*(12.8) = −2.51, *p* = .027; Table 3; Fig. 2). Responses were 6.7% faster after nap (delayed/immediate ratio = 0.933, *p* = .010) but did not change after wakefulness (ratio = 0.993, *p* = .704). The Condition main effect was not significant (*p* = .763). Accuracy increased over time (*OR* = 2.14, *p* < .001), with no Condition × Time interaction (*OR* = 0.81, *p* = .401) or Condition main effect (*p* = .315).

**Table 3.** Fixed effects of the trial-level mixed models (N = 15)

| **Term** | ***b*** | ***SE*** | ***t / z*** | ***df*** | ***p*** | ***OR*** |
| --- | --- | --- | --- | --- | --- | --- |
| ***log RT, correct trials (linear mixed model; 5529 trials)*** |  |  |  |  |  |  |
| Intercept | 6.783 | 0.032 | 211.59 | 15.5 | < .001 | — |
| Condition (nap vs wake) | 0.009 | 0.028 | 0.31 | 20.3 | .763 | — |
| Time (delayed vs immediate) | −0.038 | 0.017 | −2.28 | 13.2 | .040 | — |
| Condition × Time | −0.063 | 0.025 | −2.51 | 12.8 | .027 | — |
| ***Accuracy (logistic mixed model; 5900 trials)*** |  |  |  |  |  |  |
| Intercept | 3.726 | 0.218 | 17.09 | — | < .001 | 41.51 |
| Condition (nap vs wake) | −0.267 | 0.266 | −1.00 | — | .315 | 0.77 |
| Time (delayed vs immediate) | 0.762 | 0.171 | 4.45 | — | < .001 | 2.14 |
| Condition × Time | −0.207 | 0.246 | −0.84 | — | .401 | 0.81 |

*Note.* Effect coding: nap = +0.5, wake = −0.5; delayed test = +0.5, immediate test = −0.5. Random effects: (1 + Condition × Time | Participant) + (1 | Item) for RT; (1 + Time | Participant) + (1 | Item) for accuracy (the maximal structure was singular). df used the Satterthwaite approximation. OR = odds ratio. RT was the focal outcome for this exploratory reanalysis; accuracy was complementary.

**Fig. 2.**
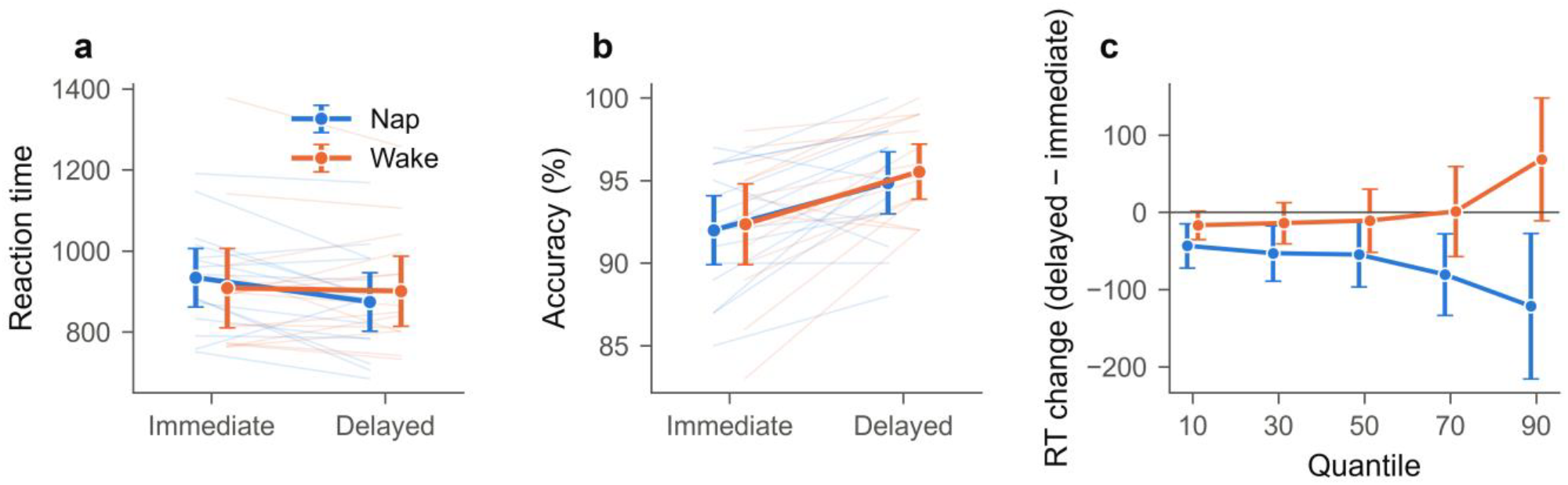
Behavioral Performance. (a) RT and (b) accuracy at immediate and delayed test (thin lines: participants; markers: means with 95% CIs). (c) Delayed − immediate RT change at the 10th–90th percentiles; negative values indicate faster delayed responses

The RT interaction increased toward the slow tail (linear trend −37.0 ms per quantile step, exact *p* = .003; Fig. 2). It ranged from −27 ms at the 10th percentile and −44 ms at the median to −190 ms at the 90th percentile (exploratory exact *p* < .001, *dz* = −1.05). The interaction was robust to alternative mean-based RT processing (exact *p* = .005–.033) and exclusion of ID07 (*b* = −0.069, *p* = .017). Median RT showed the same direction but was not significant (*p* = .111), consistent with a slow-tail contribution (Online Resource 1, Tables S1–S3).

### EEG Results

Nap showed higher delta, theta, alpha, and sigma power and lower beta power than monitored wakefulness (all Holm-corrected p ≤ .021; Fig. 3). Delta through sigma effects were robust, whereas beta was not significant with a common-average reference (Online Resource 1, Table S4).

**Fig. 3.**
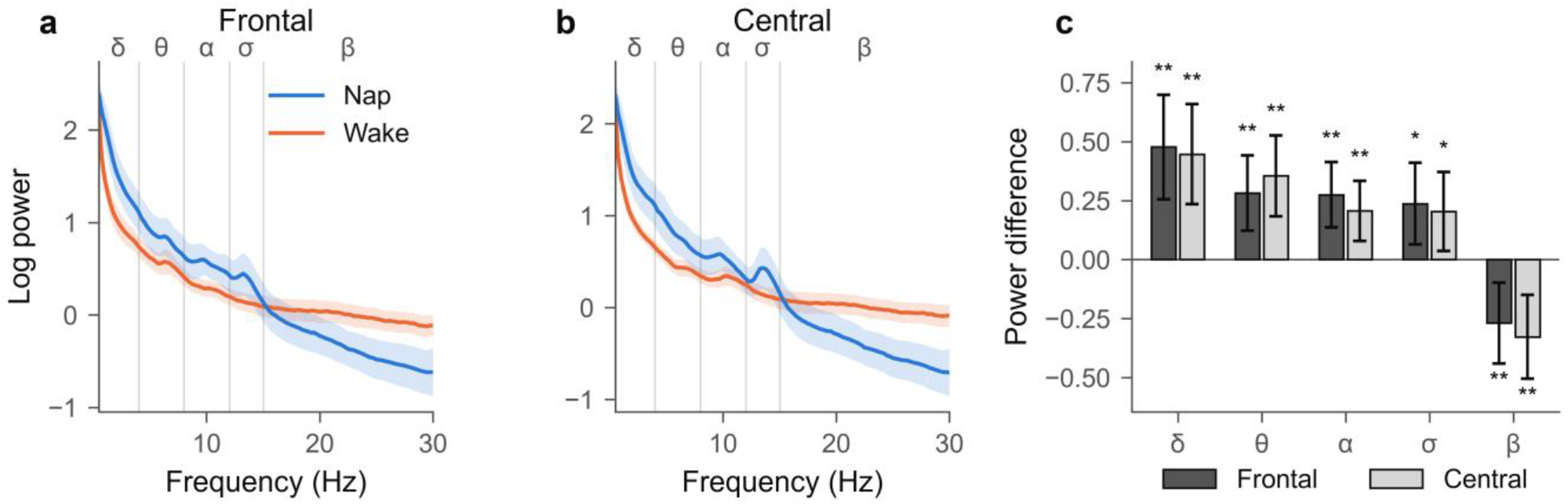
Power Spectra During the Nap and Monitored Wakefulness. (a, b) Mean log10 PSD with 95% CI at the frontal and central ROIs. (c) Mean nap − wake difference in log10 band power with 95% *CI*; Holm-corrected paired t tests, ∗*p* < .05, ∗∗*p* < .01, ∗∗∗*p* < .001

After removing the aperiodic component, periodic alpha and sigma were higher during nap (all *p* < .001; Fig. 4); periodic delta and beta did not differ. Nap had a steeper aperiodic exponent and higher offset in both regions (all *p* < .001; Fig. 5). These effects were stable across model specifications (Online Resource 1, Table S5).

**Fig. 4.**
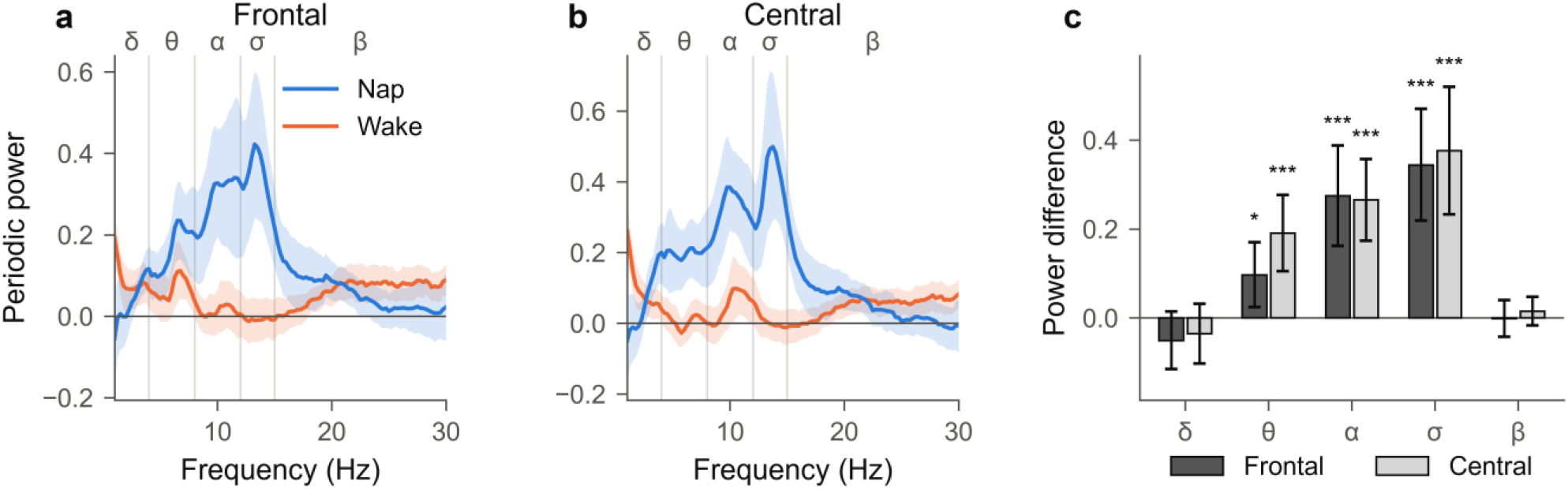
Periodic Spectral Components. (a, b) Periodic spectra after removal of the fitted aperiodic component at the frontal and central ROIs. (c) Mean nap − wake difference in periodic band power with 95% CIs; stars mark Holm-corrected significance, and non-significant contrasts are unmarked

**Fig. 5.**
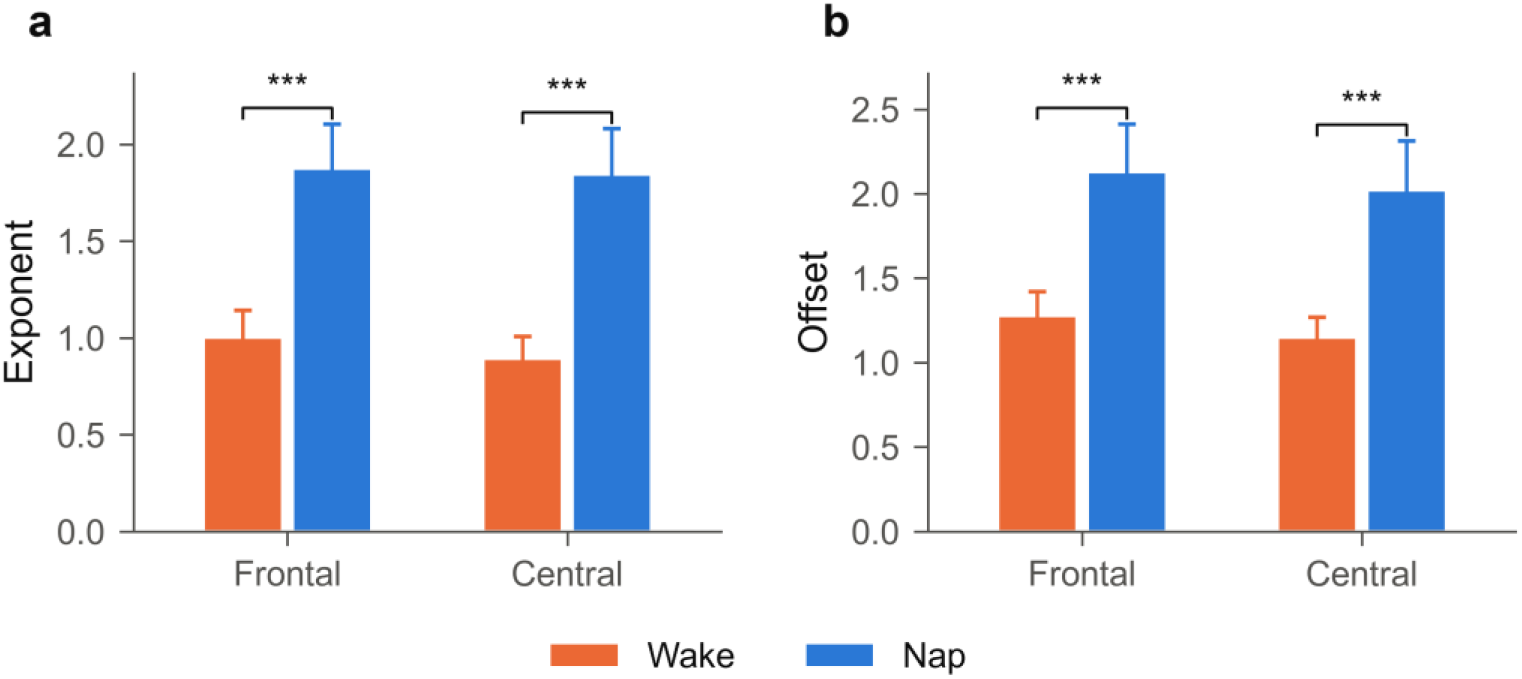
Aperiodic Spectral Parameters. (a) Aperiodic exponent and (b) offset during nap and monitored wakefulness at frontal and central ROIs. Bars show means with 95% CIs

Secondary analyses did not identify a reliable EEG correlate of the RT interaction (Online Resource 1, Tables S4– S10 and Figs. S1–S2).

## Discussion

This exploratory study found a significant Condition × Time interaction for associative-recognition RT: responses became faster after nap but not after monitored wakefulness, whereas accuracy showed no condition-specific change. The principal contribution is therefore not a broad claim that napping improved memory on every measure, but a more specific observation about the distribution of retrieval speed. The condition difference increased progressively toward the slow end of the RT distribution and was largest at the 90th percentile. This pattern is compatible with previous reports of nap-related declarative-memory benefits [3–7], while showing that a mean RT effect can conceal substantial variation across responses. A uniform shift would imply comparable speeding across most trials; the present pattern instead suggests that the nap condition was associated primarily with fewer or less extreme slow responses. Distributional localization thus provides information that neither mean RT nor accuracy alone can supply and offers a more precise behavioral target for replication. Framed this way, the potential benefit concerns response reliability as much as average speed: a condition may leave routine trials largely unchanged while reducing episodes of unusually slow retrieval. Such specificity is particularly informative when accuracy is near ceiling, because mean accuracy cannot reveal trial-to-trial instability and mean RT cannot show whether a difference is broad or confined to the upper tail.

The psychological source of the slow-tail difference remains unresolved. Prolonged responses could arise from difficult memory retrieval, transient attentional disengagement, extended evidence accumulation, response uncertainty, or delayed response selection. The present quantile analysis identifies where the condition contrast occurred but cannot distinguish among these processes. Accuracy increased from immediate to delayed test in both conditions, possibly because participants encountered the same items repeatedly and received feedback at the immediate test. High accuracy also restricted the available headroom for a selective nap effect. Importantly, faster post-nap responses were not accompanied by an observed accuracy reduction, so the RT pattern does not resemble an obvious speed–accuracy trade-off. Nevertheless, a nonsignificant accuracy interaction cannot establish equivalence between conditions. The most defensible interpretation is therefore a selective difference in retrieval speed, concentrated among the slowest responses, rather than a general enhancement of associative memory. This distinction also matters for cognitive interpretation: quicker decisions may reflect easier access to learned associations, but they could also reflect better sustained engagement or fewer transient lapses. Without independent measures of vigilance, confidence, or decision dynamics, mnemonic and attentional accounts remain observationally similar.

The EEG findings characterize the physiological contrast between the nap and monitored-wake intervals but do not establish a mechanism for the behavioral effect. Spectral parameterization was important because the conventional increase in low-frequency power did not correspond to a reliable increase in periodic delta once the aperiodic background was removed. Instead, nap was marked by a steeper aperiodic spectrum and stronger periodic alpha and sigma activity, particularly during N2. The sigma result is consistent with the prominence of spindle-range activity in non-rapid eye movement sleep [11–14], whereas the aperiodic change indicates that broadband state differences contributed substantially to the conventional band-power effects [15]. This distinction prevents the broadband low-frequency shift from being interpreted automatically as a specific delta oscillation. The stage-resolved analyses further indicated that the clearest periodic effects were expressed during established sleep, especially N2, rather than defining a single feature shared by every nap epoch. However, unequal numbers of participants contributed to the different stage estimates, N3 occurred in only four participants, and no REM sleep was scored. These stage results are therefore best treated as a physiological description of the nap opportunity rather than as evidence that one stage produced the behavioral difference. At the same time, none of the sleep-architecture, spindle, slow-wave, or spectral measures correlated reliably with behavioral change after multiplicity correction. Retrieval EEG likewise showed no nap-related interaction. The EEG results therefore verify different physiological states and refine their spectral description, but they neither mediate nor explain the RT interaction in the present sample.

Several features strengthen the analysis. The within-participant design reduces variance attributable to stable individual differences, the focal RT model retained trial-level information and item variability, and the distributional result was supported by exact permutation testing and alternative mean-based RT processing. These features are useful in a small exploratory dataset, but they do not eliminate its limitations. The sample was small and not preregistered, one immediate wake test was missing, and immediate and delayed tests reused the same items with feedback at the immediate test. Change across tests may consequently combine retention, retrieval practice, alertness, and resistance to interference. The monitored-wake interval also contained activities whose timing was not preserved, making its interval spectrum a composite of several states. A confirmatory study should preregister RT and the upper-tail contrast, recruit a larger sample, use parallel test forms or delayed-only items, preserve event markers, and measure vigilance independently. It should also test whether the slow-tail reduction replicates before using sleep or retrieval EEG to adjudicate among attentional, mnemonic, and decision-level accounts. Mechanistic claims would require a corrected and replicable association between the relevant physiological measure and behavioral change. The small sample also limits the precision of null EEG–behavior estimates, so absence of a corrected association should not be treated as evidence that sleep physiology is irrelevant. A larger design could combine RT quantiles with confidence ratings, an independent vigilance task, and trial-level retrieval EEG to test whether slow responses reflect weak memory evidence or intermittent attentional failure.

In conclusion, daytime napping was associated with faster delayed associative recognition, and the difference was concentrated in the slow tail rather than distributed uniformly across responses. Accuracy showed no condition-specific change. Nap EEG was characterized by aperiodic spectral slowing and stronger periodic sigma activity, but these measures did not explain the behavioral effect. The distributional finding narrows the behavioral phenomenon to be tested and provides a focused basis for larger confirmatory work.

## Notes

### Competing Interest Statement

The authors have declared no competing interest.

### Funding

- Ministry of Education of the People’s Republic of China: 23YJC190034
- Henan Provincial Philosophy and Social Sciences Planning Youth Special Project: 2025CJY056
- National College Students’ Innovation Training Program: 202510459037
- Henan Social Science Federation: SKL-2025-933

## References

1. Diekelmann S, Born J. The memory function of sleep. Nat Rev Neurosci. 2010;11:114–126. doi:10.1038/nrn2762

2. Stickgold R. Sleep-dependent memory consolidation. Nature. 2005;437:1272–1278. doi:10.1038/nature04286

3. Farhadian N, Khazaie H, Nami M, Khazaie S. The role of daytime napping in declarative memory performance: a systematic review. Sleep Med. 2021;84:134–141. doi:10.1016/j.sleep.2021.05.019

4. Leong RLF, Lo JC, Chee MWL. Systematic review and meta-analyses on the effects of afternoon napping on cognition. Sleep Med Rev. 2022;65:101666. doi:10.1016/j.smrv.2022.101666

5. Alger SE, Lau H, Fishbein W. Slow wave sleep during a daytime nap is necessary for protection from subsequent interference and long-term retention. Neurobiol Learn Mem. 2012;98:188–196. doi:10.1016/j.nlm.2012.06.015

6. Cousins JN, Wong KF, Raghunath BL, et al. The long-term memory benefits of a daytime nap compared with cramming. Sleep. 2019;42:zsy207. doi:10.1093/sleep/zsy207

7. Tucker MA, Hirota Y, Wamsley EJ, et al. A daytime nap containing solely non-REM sleep enhances declarative but not procedural memory. Neurobiol Learn Mem. 2006;86:241–247. doi:10.1016/j.nlm.2006.03.005

8. Qian L, Ru T, He M, et al. Effects of a brief afternoon nap on declarative and procedural memory. Neurobiol Learn Mem. 2022;194:107662. doi:10.1016/j.nlm.2022.107662

9. Lo JC, Dijk DJ, Groeger JA. Comparing the effects of nocturnal sleep and daytime napping on declarative memory consolidation. PLoS One. 2014;9:e108100. doi:10.1371/journal.pone.0108100

10. Tucker MA, Fishbein W. Enhancement of declarative memory performance following a daytime nap is contingent on strength of initial task acquisition. Sleep. 2008;31:197–203. doi:10.1093/sleep/31.2.197

11. Klinzing JG, Niethard N, Born J. Mechanisms of systems memory consolidation during sleep. Nat Neurosci. 2019;22:1598–1610. doi:10.1038/s41593-019-0467-3

12. Mölle M, Bergmann TO, Marshall L, Born J. Fast and slow spindles during the sleep slow oscillation: disparate coalescence and engagement in memory processing. Sleep. 2011;34:1411–1421. doi:10.5665/SLEEP.1290

13. Staresina BP, Bergmann TO, Bonnefond M, et al. Hierarchical nesting of slow oscillations, spindles and ripples in the human hippocampus during sleep. Nat Neurosci. 2015;18:1679–1686. doi:10.1038/nn.4119

14. Tamminen J, Lambon Ralph MA, Lewis PA. The role of sleep spindles and slow-wave activity in integrating new information in semantic memory. J Neurosci. 2013;33:15376–15381. doi:10.1523/JNEUROSCI.1333-13.2013

15. Donoghue T, Haller M, Peterson EJ, et al. Parameterizing neural power spectra into periodic and aperiodic components. Nat Neurosci. 2020;23:1655–1665. doi:10.1038/s41593-020-00744-x

16. Rugg MD, Curran T. Event-related potentials and recognition memory. Trends Cogn Sci. 2007;11:251–257. doi:10.1016/j.tics.2007.04.004

17. Hanslmayr S, Staudigl T, Fellner MC. Oscillatory power decreases and long-term memory: the information via desynchronization hypothesis. Front Hum Neurosci. 2012;6:74. doi:10.3389/fnhum.2012.00074

18. Klimesch W. EEG alpha and theta oscillations reflect cognitive and memory performance: a review and analysis. Brain Res Rev. 1999;29:169–195. doi:10.1016/S0165-0173(98)00056-3

19. Buysse DJ, Reynolds CF III, Monk TH, et al. The Pittsburgh Sleep Quality Index: a new instrument for psychiatric practice and research. Psychiatry Res. 1989;28:193–213. doi:10.1016/0165-1781(89)90047-4

20. Bastien CH, Vallières A, Morin CM. Validation of the Insomnia Severity Index as an outcome measure in insomnia research. Sleep Med. 2001;2:297–307. doi:10.1016/S1389-9457(00)00065-4

21. Beck AT, Steer RA, Brown GK. Manual for the Beck Depression Inventory-II. San Antonio: Psychological Corporation; 1996.

22. Beck AT, Epstein N, Brown G, Steer RA. An inventory for measuring clinical anxiety: psychometric properties. J Consult Clin Psychol. 1988;56:893–897. doi:10.1037/0022-006X.56.6.893

23. American Academy of Sleep Medicine. The AASM manual for the scoring of sleep and associated events: rules, terminology and technical specifications. Version 2.0.3. Darien: American Academy of Sleep Medicine; 2014.

24. Bates D, Mächler M, Bolker BM, Walker SC. Fitting linear mixed-effects models using lme4. J Stat Softw. 2015;67:1–48. doi:10.18637/jss.v067.i01

25. Kuznetsova A, Brockhoff PB, Christensen RHB. lmerTest package: tests in linear mixed effects models. J Stat Softw. 2017;82:1–26. doi:10.18637/jss.v082.i13

26. Lenth RV. emmeans: estimated marginal means, aka least-squares means. R package version 1.10.4. 2024. https://CRAN.R-project.org/package=emmeans

27. Gramfort A, Luessi M, Larson E, et al. MEG and EEG data analysis with MNE-Python. Front Neurosci. 2013;7:267. doi:10.3389/fnins.2013.00267

28. Pion-Tonachini L, Kreutz-Delgado K, Makeig S. ICLabel: an automated electroencephalographic independent component classifier, dataset, and website. Neuroimage. 2019;198:181–197. doi:10.1016/j.neuroimage.2019.05.026

29. Holm S. A simple sequentially rejective multiple test procedure. Scand J Stat. 1979;6:65–70.

30. Vallat R, Walker MP. An open-source, high-performance tool for automated sleep staging. eLife. 2021;10:e70092. doi:10.7554/eLife.70092

31. Maris E, Oostenveld R. Nonparametric statistical testing of EEG- and MEG-data. J Neurosci Methods. 2007;164:177–190. doi:10.1016/j.jneumeth.2007.03.024

32. Benjamini Y, Hochberg Y. Controlling the false discovery rate: a practical and powerful approach to multiple testing. J R Stat Soc Series B Stat Methodol. 1995;57:289–300. doi:10.1111/j.2517-6161.1995.tb02031.x

