## Supplemental Files for "Daytime Napping and Associative-Memory Retrieval Speed: An Exploratory Behavioral and EEG Study"

Journal: Sleep and Biological Rhythms

Minqi Yang[1], Ruohan Wang[1], Yawen Guan[1], Huimin Zhao[1], Weicong Wang[1], Liangdong Ji[1], Yiling Liu[1], Linxi Yan[3], and Chunyu Qu[2],*

[1] School of Education and Cognitive Science, Zhengzhou University, Zhengzhou 450001, China

[2] Department of Psychology, Ludwig-Maximilians-Universität München, Munich 80802, Germany

[3] School of Psychology, South China Normal University, Guangzhou, China

### Supplementary Contents

Behavior: Tables S1–S3. Interval and stage-specific EEG: Tables S4–S7 and Figure S1. Retrieval-phase EEG: Table S8 and Figure S2. Sleep events and sleep–behavior associations: Tables S9–S10.

### Supplementary Methods

The experimental sequence was identical in both conditions: study of 65 word pairs, a 100-trial immediate recognition test with feedback, a 60-min nap or monitored-wake interval, a 15-min post-interval delay, and a 100-trial delayed recognition test of the same items without corrective feedback. No additional learning or restudy occurred before the delayed test.

Behavioral sensitivity analyses varied RT trimming and summary rules, excluded the participant with a missing test, refitted the model after leaving out each participant, tested moderation by pair type, and compared matched-trial models with and without prestimulus posterior alpha.

Nap recordings were visually scored in consecutive 30-s epochs according to AASM criteria using F3, F4, C3, C4, O1, and O2 referenced to the contralateral mastoid together with EOG and chin EMG; no REM sleep was scored. Secondary interval-EEG analyses recomputed conventional and parameterized spectra by sleep stage, tested channel-by-frequency effects with exact cluster permutation, and detected N2 spindles and N2/N3 slow waves with YASA. Spearman associations linked sleep architecture and oscillatory measures to behavioral change, with false-discovery-rate correction across the full correlation family.

Retrieval EEG was analysed in the available immediate and delayed test recordings. Prespecified ERP summaries covered the frontal FN400 (300–500 ms) and posterior LPC (500–800 ms), with complementary whole-scalp cluster tests. Morlet-wavelet analyses summarized frontal-midline theta and posterior alpha and beta. Trial-level analyses tested whether posterior voltage predicted RT while excluding samples close to the keypress.

#### Table S1

*Participant-level behavioral robustness analyses*

| Measure | Rule | *n* | Δ*M* | 95% CI lower | 95% CI upper | Dz | *t* | p (t test) | Exact *p* | Favoring nap (n) |
|---|---|---|---|---|---|---|---|---|---|---|
| RT change, nap - wake (ms) | mean, +/-3 *SD* (primary) | 14 | -67.176 | -119.107 | -18.926 | -0.676 | -2.529 | 0.025 | 0.024 | 12 |
| RT speed-up, nap - wake (proportion) | mean, +/-3 *SD* (primary) | 14 | 0.074 | 0.025 | 0.125 | 0.74 | 2.768 | 0.016 | 0.014 | 12 |
| RT change, nap - wake (ms) | mean, +/-2.5 *SD* | 14 | -73.448 | -117.901 | -31.892 | -0.858 | -3.21 | 0.007 | 0.005 | 10 |
| RT speed-up, nap - wake (proportion) | mean, +/-2.5 *SD* | 14 | 0.081 | 0.038 | 0.128 | 0.906 | 3.39 | 0.005 | 0.003 | 10 |
| RT change, nap - wake (ms) | mean, untrimmed | 14 | -66.068 | -117.168 | -17.995 | -0.676 | -2.531 | 0.025 | 0.028 | 9 |
| RT speed-up, nap - wake (proportion) | mean, untrimmed | 14 | 0.069 | 0.02 | 0.12 | 0.704 | 2.634 | 0.021 | 0.022 | 9 |
| RT change, nap - wake (ms) | mean, 3 MAD trim | 14 | -68.043 | -120.484 | -14.271 | -0.645 | -2.414 | 0.031 | 0.033 | 11 |
| RT speed-up, nap - wake (proportion) | mean, 3 MAD trim | 14 | 0.079 | 0.024 | 0.132 | 0.732 | 2.738 | 0.017 | 0.018 | 11 |
| RT change, nap - wake (ms) | mean, 200-3000 ms | 14 | -69.033 | -112.332 | -28.509 | -0.824 | -3.083 | 0.009 | 0.007 | 10 |
| RT speed-up, nap | mean, 200-3000 | 14 | 0.073 | 0.031 | 0.118 | 0.834 | 3.122 | 0.008 | 0.006 | 11 |

| Measure | Rule | *n* | Δ*M* | 95% CI lower | 95% CI upper | Dz | *t* | p (t test) | Exact *p* | Favoring nap (n) |
|---|---|---|---|---|---|---|---|---|---|---|
| - wake (proportion) | ms | | | | | | | | | |
| RT change, nap - wake (ms) | median | 14 | -47.571 | -98.715 | 6.215 | -0.456 | -1.705 | 0.112 | 0.111 | 10 |
| RT speed-up, nap - wake (proportion) | median | 14 | 0.056 | -0.002 | 0.112 | 0.494 | 1.848 | 0.087 | 0.089 | 10 |
| RT change, nap - wake (ms) | geometric mean (log RT) | 14 | -61.572 | -105.031 | -19.621 | -0.724 | -2.708 | 0.018 | 0.018 | 10 |
| RT speed-up, nap - wake (proportion) | geometric mean (log RT) | 14 | 0.067 | 0.022 | 0.115 | 0.746 | 2.792 | 0.015 | 0.014 | 11 |
| ACC change, nap - wake | mean, +/-3 *SD* (primary) | 14 | -0.004 | -0.021 | 0.018 | -0.091 | -0.341 | 0.738 | 0.788 | 4 |
| IES change, nap - wake (ms) | mean, +/-3 *SD* (primary) | 14 | -66.92 | -131.468 | -6.88 | -0.533 | -1.993 | 0.068 | 0.072 | 10 |
| RT change, nap - wake (log RT, trial-level LMM) | (1 + cond * time_c \| subject) + (1 \| item) | 15 | -0.063 | | | | -2.507 | 0.027 | | |

*Note.* Mean differences are nap minus wake change scores. Exact p values enumerate all sign flips.

### Table S2

*RT-quantile interaction contrasts*

| Quantile | *n* | Change Nap Ms | Change Wake Ms | Interaction Ms | Dz | *t* | p (t test) | Exact *p* |
|---|---|---|---|---|---|---|---|---|
| 0.1 | 14 | -43.4 | -16.743 | -26.657 | -0.511 | -1.911 | 0.078 | 0.08 |
| 0.3 | 14 | -53.021 | -13.929 | -39.093 | -0.531 | -1.988 | 0.068 | 0.064 |
| 0.5 | 14 | -54.786 | -10.786 | -44 | -0.417 | -1.559 | 0.143 | 0.145 |
| 0.7 | 14 | -80.564 | 1.15 | -81.714 | -0.569 | -2.129 | 0.053 | 0.055 |
| 0.9 | 14 | -121.457 | 68.679 | -190.136 | -1.046 | -3.913 | 0.002 | < .001 |
| slope per quantile step | 14 | | | -36.958 | -0.936 | -3.501 | 0.004 | 0.003 |

*Note.* The contrast is (delayed − immediate)nap − (delayed − immediate)wake. Exact permutation tests used all 2^14 sign assignments. The linear trend tests whether the contrast increased toward the slow tail; individual quantiles localize that pattern and are exploratory.

### Table S3

*Key sensitivity analyses for log RT*

| Analysis | Subset | Term | Estimate | *SE* | *df* | *t* | *p* | Observations | Participants | Singular |
|---|---|---|---|---|---|---|---|---|---|---|
| pair-type model | moderation | cond:time_c:pair | 0.029 | 0.03 | 5270.985 | 0.955 | 0.34 | 5529 | 15 | False |
| pair-type simple interaction | recombined | cond:time_c | -0.081 | 0.032 | 34.401 | -2.55 | 0.015 | 5529 | 15 | False |
| pair-type simple interaction | intact | cond:time_c | -0.053 | 0.027 | 17.375 | -1.957 | 0.067 | 5529 | 15 | False |
| focal participant-slope structure | without alpha | cond:time_c | -0.068 | 0.027 | 10.092 | -2.517 | 0.03 | 4987 | 15 | False |
| focal participant-slope structure | with alpha | cond:time_c | -0.068 | 0.027 | 10.088 | -2.523 | 0.03 | 4987 | 15 | False |

*Note.* This compact table retains the sensitivity analyses that materially inform interpretation: pair-type moderation and matched-trial models with and without prestimulus alpha.

Prestimulus alpha was estimated at 8–12 Hz over P3, Pz, P4, PO3, POz, and PO4 from −700 to −300 ms, log10-transformed, and standardized within session before entry into the matched-trial model.

Leave-one-participant-out summary: all 15 interaction estimates were negative ($b$ = -0.072 to -0.050); p values ranged from 0.015 to 0.056, and 12 of 15 fits had $p < .05$. No exclusion reversed the effect direction.

**Table S4**

*Summary of conventional band-power robustness analyses*

| ROI | Band | Variants | Min Dz | Max Dz | Largest P Holm | Significant Variants |
|---|---|---|---|---|---|---|
| central | alpha | 7 | 0.831 | 1.421 | 0.017 | 7 |
| central | beta | 7 | -1.386 | -0.305 | 0.257 | 6 |
| central | delta | 7 | 1.103 | 1.782 | 0.005 | 7 |
| central | sigma | 7 | 0.618 | 1.113 | 0.038 | 7 |
| central | theta | 7 | 1.081 | 2.789 | 0.005 | 7 |
| frontal | alpha | 7 | 1.031 | 1.286 | 0.006 | 7 |
| frontal | beta | 7 | -1.159 | -0.44 | 0.111 | 6 |
| frontal | delta | 7 | 1.124 | 1.545 | 0.004 | 7 |
| frontal | sigma | 7 | 0.707 | 1.055 | 0.02 | 7 |
| frontal | theta | 7 | 0.912 | 2.241 | 0.009 | 7 |

*Note.* The seven processing variants covered artefact thresholds, muscle rejection, reference scheme, participant exclusion, and duration matching. The table reports the range of effect sizes and the largest Holm-corrected p value rather than repeating every model row.

**Table S5**

*Summary of spectral-parameterization sensitivity*

| ROI | Measure | Specifications | Min Dz | Max Dz | Significant Specifications |
|---|---|---|---|---|---|
| central | exponent | 6 | 1.899 | 2.383 | 6 |
| central | offset | 6 | 1.807 | 2.392 | 6 |
| central | periodic_alpha | 6 | 0.553 | 1.598 | 5 |
| central | periodic_beta | 6 | -0.031 | 0.356 | 0 |
| central | periodic_delta | 6 | -0.67 | 0.057 | 1 |
| central | periodic_sigma | 6 | 1.021 | 1.453 | 6 |
| central | periodic_theta | 6 | -0.395 | 1.227 | 3 |
| frontal | exponent | 6 | 1.559 | 2.117 | 6 |
| frontal | offset | 6 | 1.521 | 2.151 | 6 |
| frontal | periodic_alpha | 6 | 0.79 | 1.349 | 6 |
| frontal | periodic_beta | 6 | -0.176 | 0.042 | 0 |
| frontal | periodic_delta | 6 | -1.18 | 0.031 | 2 |
| frontal | periodic_sigma | 6 | 1.181 | 1.519 | 6 |
| frontal | periodic_theta | 6 | -1.017 | 0.734 | 5 |

*Note.* Rows summarize six combinations of model mode and fitting range. Counts are descriptive because the specifications are alternative analyses of the same data.

**Table S6**

*Significant channel × frequency clusters*

| Cluster | Sign | *p* | T Sum | Freq Min | Freq Max | N Channels | N Points |
|---|---|---|---|---|---|---|---|
| 1 | nap < wake | < .001 | -17492.018 | 7.5 | 30 | 57 | 3372 |
| 0 | nap > wake | 0.025 | 7278.29 | 0.5 | 15 | 60 | 1993 |
| 2 | nap < wake | 0.884 | -2.159 | 12.25 | 12.25 | 1 | 1 |

*Note.* Cluster p values are exact sign-flip probabilities; inference applies to each cluster as a whole.

**Table S7**

*Stage-Specific Spectral Analyses*

**Panel A.** *Data availability*

| State | Participants | Median Minutes | Minimum Minutes | Maximum Minutes |
|---|---|---|---|---|
| N1 | 15 | 9.5 | 2.5 | 17 |
| N2 | 13 | 29.5 | 12 | 51.5 |
| N3 | 3 | 7 | 3 | 18.5 |
| nap-period wake | 13 | 12.5 | 3.5 | 46 |
| monitored wake | 15 | 58 | 30 | 61 |

*Note.* A state was included for a participant when at least five clean 30-s epochs were available. Values summarize included participants rather than listing every participant-state cell.

**Panel B.** *Conventional band-power contrasts*

| State | ROI | Measure | *n* | Stage *M* | Wake *M* | Δ*M* | Dz | *t* | Holm *p* | Exact Holm *p* |
|---|---|---|---|---|---|---|---|---|---|---|
| N2 | frontal | Delta | 13 | 2.53 | 1.845 | 0.685 | 2.009 | 7.244 | < .001 | 0.002 |
| N3 | frontal | Delta | 3 | 3.162 | 1.738 | 1.423 | 23.576 | 40.835 | 0.003 | 1 |
| N2 | central | Delta | 13 | 2.447 | 1.783 | 0.663 | 2.541 | 9.162 | < .001 | 0.001 |
| N3 | central | Delta | 3 | 3.108 | 1.732 | 1.376 | 12.455 | 21.573 | 0.011 | 1 |
| N2 | frontal | Theta | 13 | 1.694 | 1.214 | 0.479 | 1.835 | 6.618 | < .001 | 0.002 |
| N3 | frontal | Theta | 3 | 1.891 | 1.214 | 0.677 | 4.044 | 7.005 | 0.059 | 1 |
| N2 | central | Theta | 13 | 1.684 | 1.107 | 0.576 | 2.25 | 8.111 | < .001 | 0.001 |
| N3 | central | Theta | 3 | 1.789 | 1.145 | 0.644 | 3.38 | 5.855 | 0.084 | 1 |
| N2 | frontal | Alpha | 13 | 1.236 | 0.925 | 0.311 | 1.526 | 5.5 | < .001 | 0.002 |
| N3 | frontal | Alpha | 3 | 1.259 | 0.878 | 0.381 | 1.879 | 3.255 | 0.166 | 1 |
| N2 | central | Alpha | 13 | 1.158 | 0.95 | 0.208 | 1.153 | 4.156 | 0.001 | 0.003 |
| N3 | central | Alpha | 3 | 1.072 | 0.912 | 0.16 | 1.558 | 2.698 | 0.229 | 1 |
| N2 | frontal | Sigma | 13 | 1.09 | 0.641 | 0.449 | 1.93 | 6.96 | < .001 | 0.001 |
| N3 | frontal | Sigma | 3 | 0.823 | 0.556 | 0.267 | 0.985 | 1.707 | 0.23 | 1 |
| N2 | central | Sigma | 13 | 1.074 | 0.654 | 0.419 | 2.061 | 7.43 | < .001 | 0.001 |
| N3 | central | Sigma | 3 | 0.791 | 0.553 | 0.238 | 1.268 | 2.197 | 0.229 | 1 |
| N2 | frontal | Beta | 13 | 0.852 | 1.182 | -0.33 | -1.323 | -4.768 | < .001 | 0.002 |
| N3 | frontal | Beta | 3 | 0.447 | 1.134 | -0.687 | -13.61 | -23.574 | 0.007 | 1 |
| N2 | central | Beta | 13 | 0.806 | 1.186 | -0.38 | -1.474 | -5.316 | < .001 | 0.001 |
| N3 | central | Beta | 3 | 0.464 | 1.088 | -0.624 | -5.504 | -9.534 | 0.043 | 1 |

*Note.* Power values are log10-integrated power. Holm correction was applied across frequency bands within each state and ROI; exact p values are from complete sign-flip enumeration. N3 estimates are descriptive because only three participants met the duration rule.

**Panel C.** *Aperiodic and periodic spectral contrasts*

| Family | State | ROI | Measure | *n* | Stage *M* | Wake *M* | Δ*M* | Dz | *t* | Holm *p* | Exact Holm *p* |
|---|---|---|---|---|---|---|---|---|---|---|---|
| Aperiodic | N2 | frontal | Offset | 13 | 2.523 | 1.297 | 1.226 | 3.142 | 11.328 | < .001 | < .001 |
| Aperiodic | N2 | frontal | Exponent | 13 | 2.227 | 1.01 | 1.217 | 3.686 | 13.289 | < .001 | < .001 |
| Aperiodic | N3 | frontal | Offset | 3 | 3.314 | 1.269 | 2.045 | 19.884 | 34.441 | < .001 | 0.5 |
| Aperiodic | N3 | frontal | Exponent | 3 | 3.083 | 1.02 | 2.063 | 26.14 | 45.276 | < .001 | 0.5 |
| Aperiodic | N2 | central | Offset | 13 | 2.421 | 1.177 | 1.245 | 4.45 | 16.045 | < .001 | < .001 |
| Aperiodic | N2 | central | Exponent | 13 | 2.209 | 0.904 | 1.305 | 5.953 | 21.465 | < .001 | < .001 |
| Aperiodic | N3 | central | Offset | 3 | 3.207 | 1.277 | 1.931 | 11.069 | 19.172 | 0.005 | 0.5 |
| Aperiodic | N3 | central | Exponent | 3 | 3.015 | 1.052 | 1.963 | 10.946 | 18.958 | 0.005 | 0.5 |
| Periodic | N2 | frontal | Delta | 13 | 0.04 | 0.087 | -0.047 | -0.456 | -1.644 | 0.252 | 0.257 |

| **Family** | **State** | **ROI** | **Measure** | ***n*** | **Stage *M*** | **Wake *M*** | **Δ*M*** | **Dz** | ***t*** | **Holm *p*** | **Exact Holm *p*** |
|---|---|---|---|---|---|---|---|---|---|---|---|
| Periodic | N2 | frontal | Theta | 13 | 0.252 | 0.076 | 0.175 | 1.078 | 3.886 | 0.006 | 0.007 |
| Periodic | N2 | frontal | Alpha | 13 | 0.316 | 0.027 | 0.289 | 1.427 | 5.144 | < .001 | 0.002 |
| Periodic | N2 | frontal | Sigma | 13 | 0.566 | 0.005 | 0.56 | 3.084 | 11.12 | < .001 | 0.001 |
| Periodic | N2 | frontal | Beta | 13 | 0.045 | 0.057 | -0.012 | -0.217 | -0.783 | 0.449 | 0.457 |
| Periodic | N3 | frontal | Delta | 3 | 0.016 | 0.074 | -0.059 | -1.055 | -1.828 | 0.418 | 1 |
| Periodic | N3 | frontal | Theta | 3 | 0.304 | 0.11 | 0.194 | 1.47 | 2.546 | 0.377 | 1 |
| Periodic | N3 | frontal | Alpha | 3 | 0.407 | 0.019 | 0.387 | 2.426 | 4.201 | 0.209 | 1 |
| Periodic | N3 | frontal | Sigma | 3 | 0.456 | -0.04 | 0.496 | 2.989 | 5.177 | 0.177 | 1 |
| Periodic | N3 | frontal | Beta | 3 | 0.004 | 0.051 | -0.047 | -0.831 | -1.439 | 0.418 | 1 |
| Periodic | N2 | central | Delta | 13 | 0.064 | 0.09 | -0.026 | -0.267 | -0.963 | 0.709 | 0.718 |
| Periodic | N2 | central | Theta | 13 | 0.309 | 0.014 | 0.296 | 1.824 | 6.575 | < .001 | 0.001 |
| Periodic | N2 | central | Alpha | 13 | 0.321 | 0.065 | 0.256 | 1.733 | 6.247 | < .001 | 0.001 |
| Periodic | N2 | central | Sigma | 13 | 0.618 | 0.018 | 0.6 | 2.958 | 10.664 | < .001 | 0.001 |
| Periodic | N2 | central | Beta | 13 | 0.05 | 0.04 | 0.009 | 0.17 | 0.614 | 0.709 | 0.718 |
| Periodic | N3 | central | Delta | 3 | 0.031 | 0.064 | -0.033 | -0.409 | -0.708 | 0.69 | 1 |
| Periodic | N3 | central | Theta | 3 | 0.242 | 0.061 | 0.182 | 1.557 | 2.697 | 0.343 | 1 |
| Periodic | N3 | central | Alpha | 3 | 0.245 | 0.075 | 0.169 | 3.588 | 6.215 | 0.125 | 1 |
| Periodic | N3 | central | Sigma | 3 | 0.477 | -0.018 | 0.495 | 3.311 | 5.736 | 0.125 | 1 |
| Periodic | N3 | central | Beta | 3 | 0.016 | 0.041 | -0.025 | -0.708 | -1.227 | 0.69 | 1 |

*Note.* Spectra were parameterized in fixed mode over 1–30 Hz; fits with $R^2 \leq .75$ were excluded. Holm correction was applied within each state, ROI, and parameter family. N3 estimates are descriptive because only three participants met the duration rule.

**Figure S1**

*Stage-Specific Spectra and N2/N3 Contrasts Against Independent Monitored Wakefulness*

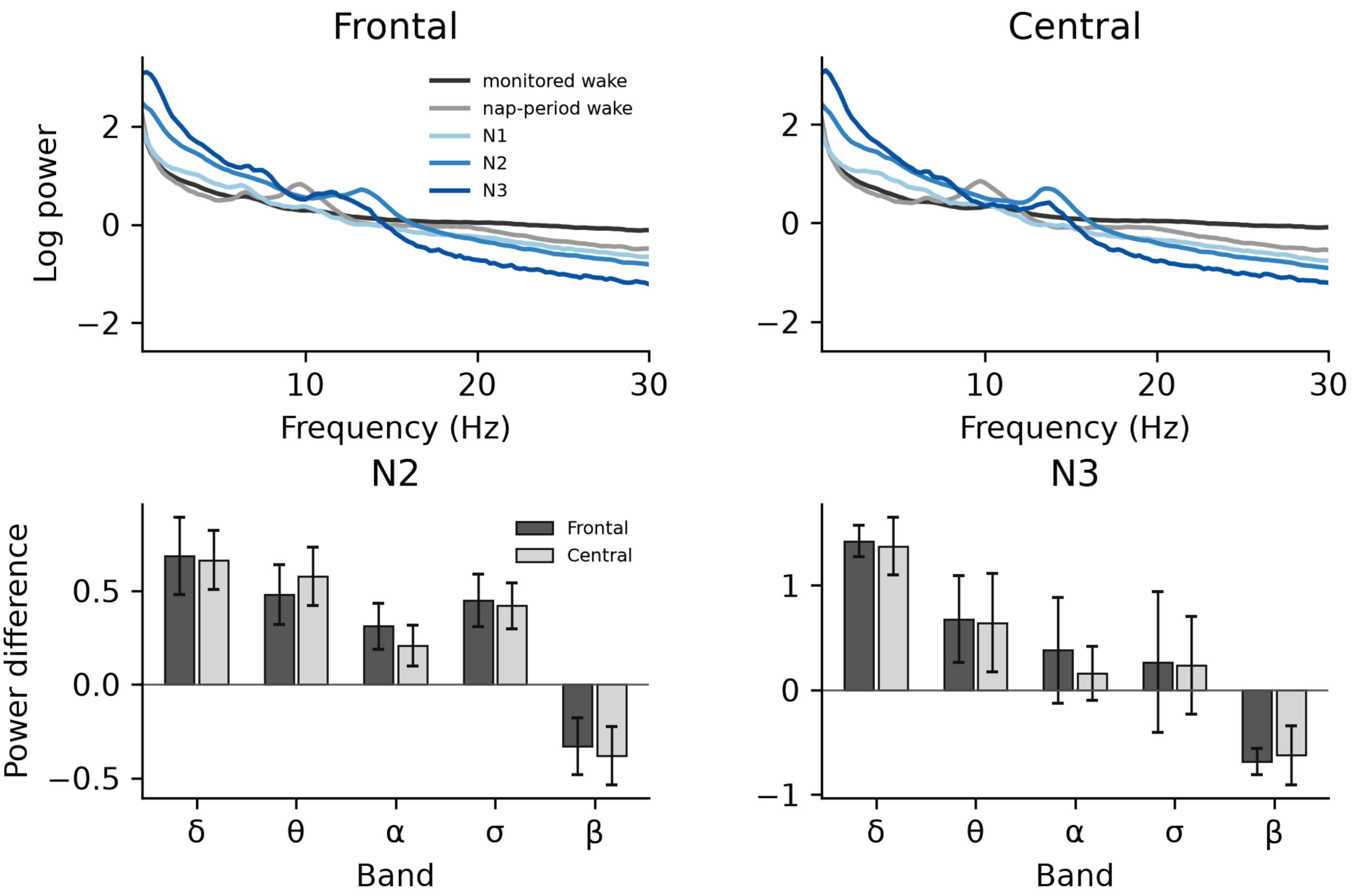


*Note.* Lines show state means; bars show mean paired differences with 95% confidence intervals. Sample sizes are reported in Table S7, Panel A; N3 contrasts are descriptive (n = 3).

**Table S8**

*Secondary Retrieval-Phase EEG Analyses*

**Panel A.** *Key trial-level retrieval-ERP contrasts*

| Component | Term | Estimate | Std. Error | *df* | T Value | Pr(>\|T\|) | Holm *p* | Observations | Participants |
|---|---|---|---|---|---|---|---|---|---|
| FN400 | pair | 1.445 | 0.392 | 4927.402 | 3.687 | < .001 | < .001 | 4987 | 15 |
| FN400 | cond:time_c | -3.064 | 1.899 | 39.859 | -1.613 | 0.115 | 0.229 | 4987 | 15 |
| FN400 | cond:time_c:pair | -0.069 | 1.568 | 4927.438 | -0.044 | 0.965 | 0.965 | 4987 | 15 |
| LPC | pair | 1.723 | 0.45 | 4933.712 | 3.829 | < .001 | < .001 | 4987 | 15 |
| LPC | cond:time_c | -1.268 | 1.156 | 44.419 | -1.097 | 0.279 | 0.279 | 4987 | 15 |
| LPC | cond:time_c:pair | 3.227 | 1.8 | 4934.161 | 1.793 | 0.073 | 0.146 | 4987 | 15 |

*Note.* Only the old/new, Condition × Time, and three-way contrasts are retained. Models included participant, recording-session, and item random intercepts; Holm correction was applied across components.

**Panel B.** *Minimum-p spatiotemporal ERP cluster for each planned contrast*

| Contrast | *n* | Sign | *p* | T Sum | T Start Ms | T End Ms | N Channels |
|---|---|---|---|---|---|---|---|
| cond_x_time | 13 | - | 0.146 | -1119.745 | 385 | 550 | 35 |
| cond_x_time_x_pair | 13 | + | 0.734 | 92.66 | 685 | 725 | 10 |
| old_new | 13 | + | 0.021 | 7139.183 | 380 | 830 | 57 |

*Note.* Cluster inference covered all scalp channels and 0–1,000 ms. Only the old/new contrast reached significance; the minimum-p cluster is shown for each contrast to summarize the null nap-related tests without listing every supra-threshold fragment.

**Panel C.** *Retrieval-phase time-frequency contrasts*

| ROI | Contrast | *n* | Mean Db | Dz | *t* | *p* | Exact *p* | Holm *p* |
|---|---|---|---|---|---|---|---|---|
| theta_fm | old_new | 13 | 0.123 | 0.197 | 0.712 | 0.49 | 0.482 | 0.709 |
| theta_fm | cond_x_time | 13 | -0.571 | -0.426 | -1.535 | 0.151 | 0.164 | 0.492 |
| theta_fm | cond_x_time_x_pair | 13 | -0.312 | -0.156 | -0.563 | 0.584 | 0.574 | 1 |
| theta_fm | task_vs_baseline | 13 | 0.879 | 0.698 | 2.518 | 0.027 | 0.029 | 0.029 |
| alpha_post | old_new | 13 | 0.269 | 0.542 | 1.953 | 0.075 | 0.073 | 0.22 |
| alpha_post | cond_x_time | 13 | 0.383 | 0.211 | 0.762 | 0.461 | 0.473 | 0.55 |
| alpha_post | cond_x_time_x_pair | 13 | -0.644 | -0.257 | -0.926 | 0.373 | 0.365 | 1 |
| alpha_post | task_vs_baseline | 13 | -1.928 | -0.688 | -2.479 | 0.029 | 0.006 | 0.012 |
| beta_post | old_new | 13 | 0.142 | 0.265 | 0.957 | 0.358 | 0.355 | 0.709 |
| beta_post | cond_x_time | 13 | 0.327 | 0.311 | 1.122 | 0.284 | 0.275 | 0.55 |
| beta_post | cond_x_time_x_pair | 13 | 0.115 | 0.069 | 0.248 | 0.808 | 0.799 | 1 |
| beta_post | task_vs_baseline | 13 | -1.81 | -1.462 | -5.271 | < .001 | < .001 | < .001 |

*Note.* Power was expressed in dB relative to the prestimulus baseline. Exact sign-flip p values were Holm-corrected across the three prespecified ROIs within each contrast family.

**Panel D.** *Significant time-resolved within-participant ERP–RT clusters*

| ROI | Start Ms | End Ms | Sign | Mean R | *p* | *n* |
|---|---|---|---|---|---|---|
| FN400 | 485 | 665 | - | -0.071 | 0.002 | 15 |
| LPC | 445 | 825 | - | -0.109 | < .001 | 15 |

*Note.* Correlations were computed within participant before group-level cluster correction. Samples within 100 ms of the keypress were excluded.

**Figure S2**

*Retrieval-Related Time–Frequency Activity*

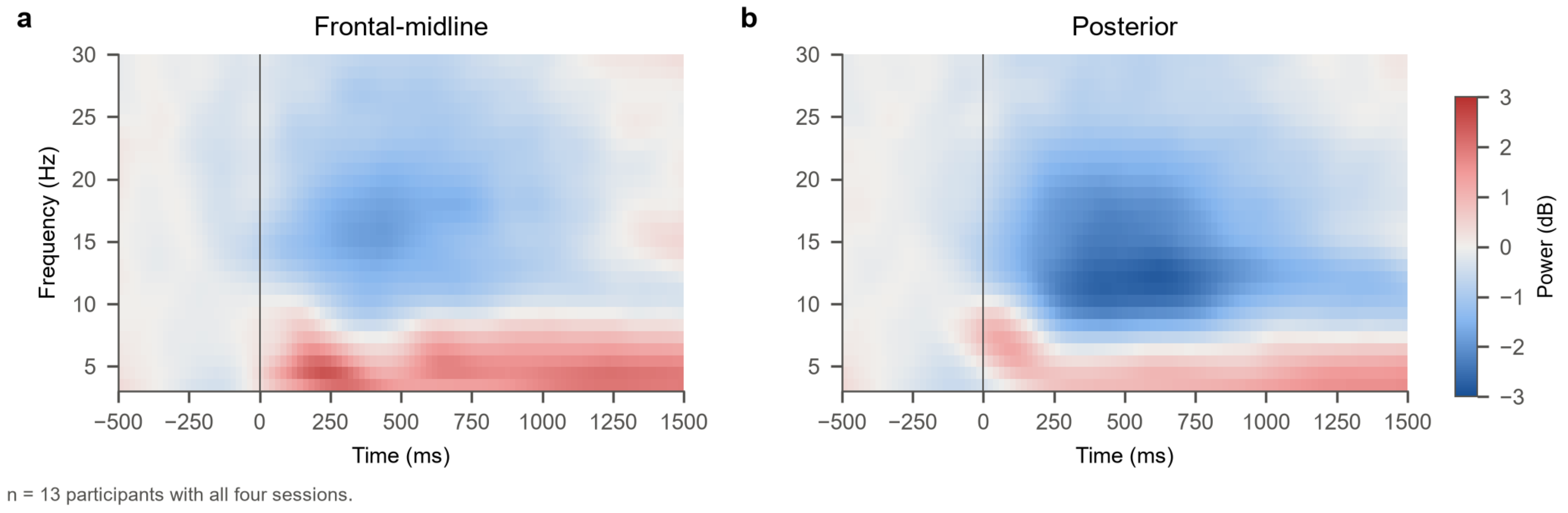


*Note.* Participant-weighted task-related power relative to the prestimulus baseline is shown at the frontal-midline and posterior ROIs for the 13 participants with all four test recordings.

**Table S9**

*Descriptive sleep-event measures*

| Measure | *n* | *M* | *SD* | Minimum | Maximum |
|---|---|---|---|---|---|
| Frontal spindle density (/min) | 13 | 1.54 | 1.05 | 0.13 | 4.14 |
| Parietal spindle density (/min) | 13 | 2.26 | 1.46 | 0.07 | 5 |
| Frontal spindle amplitude (µV) | 13 | 72.47 | 18.89 | 51.81 | 107.21 |
| Parietal spindle amplitude (µV) | 13 | 57.8 | 17.7 | 37.7 | 89.29 |
| Frontal spindle frequency (Hz) | 13 | 13.3 | 0.31 | 12.83 | 13.88 |
| Parietal spindle frequency (Hz) | 13 | 13.76 | 0.36 | 13.12 | 14.3 |
| Frontal slow-wave density (/min) | 13 | 3.92 | 4.25 | 0.41 | 12.42 |
| Frontal slow-wave peak-to-peak amplitude (µV) | 13 | 118.93 | 9.82 | 105.13 | 133.57 |

*Note.* Spindles were detected during N2 and slow waves during N2 + N3. Amplitude and frequency are defined only when at least one event was detected; zeros for density indicate that the participant had eligible sleep but no detected event.

**Table S10**

*Sleep–behavior Spearman correlations*

| Sleep measure | Behavioral outcome | *n* | ρ | *p* | FDR-adjusted *p* |
|---|---|---|---|---|---|
| TST | Accuracy change | 15 | -0.403 | 0.137 | 0.883 |
| TST | Relative RT improvement | 15 | -0.014 | 0.96 | 0.986 |
| TST | Accuracy retention | 15 | -0.452 | 0.091 | 0.883 |

| Sleep measure | Behavioral outcome | *n* | ρ | *p* | FDR-adjusted *p* |
|---|---|---|---|---|---|
| TST | IES change | 15 | 0.234 | 0.401 | 0.907 |
| N2 duration | Accuracy change | 15 | -0.296 | 0.283 | 0.883 |
| N2 duration | Relative RT improvement | 15 | 0.011 | 0.97 | 0.986 |
| N2 duration | Accuracy retention | 15 | -0.329 | 0.23 | 0.883 |
| N2 duration | IES change | 15 | 0.191 | 0.494 | 0.907 |
| N3 duration | Accuracy change | 15 | -0.29 | 0.294 | 0.883 |
| N3 duration | Relative RT improvement | 15 | -0.303 | 0.273 | 0.883 |
| N3 duration | Accuracy retention | 15 | -0.36 | 0.187 | 0.883 |
| N3 duration | IES change | 15 | 0.325 | 0.237 | 0.883 |
| Frontal spindle density | Accuracy change | 13 | 0.202 | 0.508 | 0.907 |
| Frontal spindle density | Relative RT improvement | 13 | 0.071 | 0.817 | 0.94 |
| Frontal spindle density | Accuracy retention | 13 | 0.289 | 0.338 | 0.907 |
| Frontal spindle density | IES change | 13 | -0.005 | 0.986 | 0.986 |
| Frontal spindle amplitude | Accuracy change | 13 | -0.113 | 0.712 | 0.937 |
| Frontal spindle amplitude | Relative RT improvement | 13 | 0.302 | 0.316 | 0.902 |
| Frontal spindle amplitude | Accuracy retention | 13 | -0.072 | 0.816 | 0.94 |
| Frontal spindle amplitude | IES change | 13 | -0.231 | 0.448 | 0.907 |
| Frontal spindle frequency | Accuracy change | 13 | 0.008 | 0.979 | 0.986 |
| Frontal spindle frequency | Relative RT improvement | 13 | 0.341 | 0.255 | 0.883 |
| Frontal spindle frequency | Accuracy retention | 13 | 0.121 | 0.694 | 0.937 |
| Frontal spindle frequency | IES change | 13 | -0.462 | 0.112 | 0.883 |
| Frontal slow-wave density | Accuracy change | 13 | -0.373 | 0.209 | 0.883 |
| Frontal slow-wave density | Relative RT improvement | 13 | -0.088 | 0.775 | 0.94 |
| Frontal slow-wave density | Accuracy retention | 13 | -0.413 | 0.161 | 0.883 |
| Frontal slow-wave density | IES change | 13 | 0.11 | 0.721 | 0.937 |
| Frontal slow-wave amplitude | Accuracy change | 13 | -0.343 | 0.251 | 0.883 |
| Frontal slow-wave amplitude | Relative RT improvement | 13 | 0.489 | 0.09 | 0.883 |
| Frontal slow-wave amplitude | Accuracy retention | 13 | -0.391 | 0.187 | 0.883 |
| Frontal slow-wave amplitude | IES change | 13 | -0.352 | 0.239 | 0.883 |
| Parietal spindle density | Accuracy change | 13 | -0.263 | 0.386 | 0.907 |
| Parietal spindle density | Relative RT improvement | 13 | 0.11 | 0.721 | 0.937 |
| Parietal spindle density | Accuracy retention | 13 | -0.228 | 0.453 | 0.907 |
| Parietal spindle density | IES change | 13 | 0.055 | 0.859 | 0.954 |
| Parietal spindle amplitude | Accuracy change | 13 | -0.221 | 0.467 | 0.907 |
| Parietal spindle amplitude | Relative RT improvement | 13 | -0.159 | 0.603 | 0.937 |
| Parietal spindle amplitude | Accuracy retention | 13 | -0.162 | 0.596 | 0.937 |
| Parietal spindle amplitude | IES change | 13 | 0.357 | 0.231 | 0.883 |
| Parietal spindle frequency | Accuracy change | 13 | -0.091 | 0.767 | 0.94 |
| Parietal spindle frequency | Relative RT improvement | 13 | -0.137 | 0.655 | 0.937 |
| Parietal spindle frequency | Accuracy retention | 13 | 0.039 | 0.901 | 0.982 |
| Parietal spindle frequency | IES change | 13 | 0.077 | 0.803 | 0.94 |
| Log fast-sigma power | Accuracy change | 13 | -0.315 | 0.294 | 0.883 |
| Log fast-sigma power | Relative RT improvement | 13 | -0.022 | 0.943 | 0.986 |
| Log fast-sigma power | Accuracy retention | 13 | -0.223 | 0.464 | 0.907 |
| Log fast-sigma power | IES change | 13 | 0.231 | 0.448 | 0.907 |
| Log slow-sigma power | Accuracy change | 13 | -0.199 | 0.514 | 0.907 |
| Log slow-sigma power | Relative RT improvement | 13 | 0.209 | 0.494 | 0.907 |
| Log slow-sigma power | Accuracy retention | 13 | -0.168 | 0.584 | 0.937 |

| **Sleep measure** | **Behavioral outcome** | ***n*** | **ρ** | ***p*** | **FDR-adjusted *p*** |
|---|---|---|---|---|---|
| Log slow-sigma power | IES change | 13 | -0.066 | 0.831 | 0.94 |
| Log sigma power | Accuracy change | 13 | -0.183 | 0.55 | 0.937 |
| Log sigma power | Relative RT improvement | 13 | 0.22 | 0.471 | 0.907 |
| Log sigma power | Accuracy retention | 13 | -0.129 | 0.674 | 0.937 |
| Log sigma power | IES change | 13 | -0.104 | 0.734 | 0.937 |
| Log SWA | Accuracy change | 13 | -0.318 | 0.289 | 0.883 |
| Log SWA | Relative RT improvement | 13 | -0.143 | 0.642 | 0.937 |
| Log SWA | Accuracy retention | 13 | -0.352 | 0.238 | 0.883 |
| Log SWA | IES change | 13 | 0.143 | 0.642 | 0.937 |

*Note.* All tested sleep-measure × behavioral-outcome combinations are shown. FDR p values use the Benjamini–Hochberg procedure across the full family of 60 tests; none survived correction. REM duration was not tested because no REM sleep was scored.
